# Mechanism of phosphorylation dependent interactions of complete Retinoblastoma-AB pocket domain with its linker

**DOI:** 10.1101/2025.05.27.656056

**Authors:** K. Kavana, A. Bhattacherjee, I. Ghosh, S. Tiwari

## Abstract

Retinoblastoma protein (pRb) is a critical tumor suppressor, whose activity is regulated through phosphorylation of Serine/Threonine residues within unstructured linker regions. Due to the absence of high-resolution structures encompassing these linkers, many studies have indirectly examined the phosphorylation effects on pRb-protein interactions. Here, we present computational models of the full pRb pocket domain (pRbAB), incorporating the flexible linker (pRbPL) that connects boxes A and B. Three models, generated by ROBETTA and AlphaFold, were used to explore the mechanisms by which phosphorylation at Ser608 and Ser612 influence the interaction of pRbAB with E2F transactivation domain (E2FTA). All models revealed a short helix (residues 602–607), similar to that observed in a previously published crystal structure of pRbAB with shortened pRbPL linker and phosphomimetic mutation at Ser608, with E2FTA (PDB:4ELL), and proposed to competitively inhibit E2FTA binding. Interestingly, this helix formed irrespective of phosphorylation, indicating that additional factors influencing the interaction upon phosphorylation. In our models, the pRbPL linker displayed significant motion, especially near Ser608 and Ser612, facilitating phosphorylation, but without inducing large conformational changes in the pocket domains. Phosphorylation at Ser608 and Ser612 resulted in changes in multiple parameters including electrostatic properties, compactness and energy landscape of pRbAB constituting important mechanisms that promote E2F release. Additionally, recapitulating known R661W mutation in these models correctly showed reduced E2FTA binding, demonstrating the predictive power of these models to study the effects of missense mutations on the stability, conformation and interactions of pRbAB with E2FTA.

## 1. INTRODUCTION

The retinoblastoma (pRb) tumor suppressor protein is a multifunctional, stable protein that is considered to be a factor in the development of multicellularity[1]. It is a “hub-protein” in cell-cycle interaction network, and has hundreds of interacting partners [2]. The best understood role of pRb is in the regulation of cell cycle, but it also governs decisions about differentiation, senescence, and maintenance of genomic epigenetic stability [3]. Loss of pRb and its associated pathways is considered a hallmark of cancer. Cancer genomics studies show mut tions in pRb that result in loss of its function in a wide variety of cancers, as well as mutations in pRb associated pathways in diverse cancers [4, 5]. Loss of pRb function has implications for both chemosensitivity to therapy and lineage plasticity in case of relapse.

The folded domains of pRb include a globular N-terminal domain (pRbN), the “pocket-domain (pRbAB)” comprising of A and B boxes (pRbA and pRbB), and a C-terminal domain (pRbC) (Fig. 1A). These folded domains are linked via disordered linkers (Fig. 1). Both pRbN and pRbAB are cyclin-fold domains. pRbAB forms the main interaction domain of pRb and is termed “small pocket”. pRbAB along with pRbC, forms the “large pocket”. The conserved interaction surfaces of the pRbAB are the E2F transcription factor binding site at the highly conserved interface between the A- and B-box [6] and the LxCxE motif binding pocket in the B-box (Fig. 1B) [7]. The LxCxE motif, is present in many pRb-interacting proteins and in some transforming viral proteins. The viral proteins containing this motif, e.g., E7 protein of human papilloma virus can displace E2F by binding to pRb via a high affinity LxCxE based interactions [8]. Association of pRb with the cell cycle promoting E2F transcription factors is dependent on the phosphorylation status of pRb [9]. It has been proposed that pRb uses both E2F- and LxCxE-dependent, and independent pathways parallelly to suppress tumorigenesis [10]. Therefore, pRb and its associated pathways are considered important targets to prevent aberrant proliferation.

**Fig. 1:**
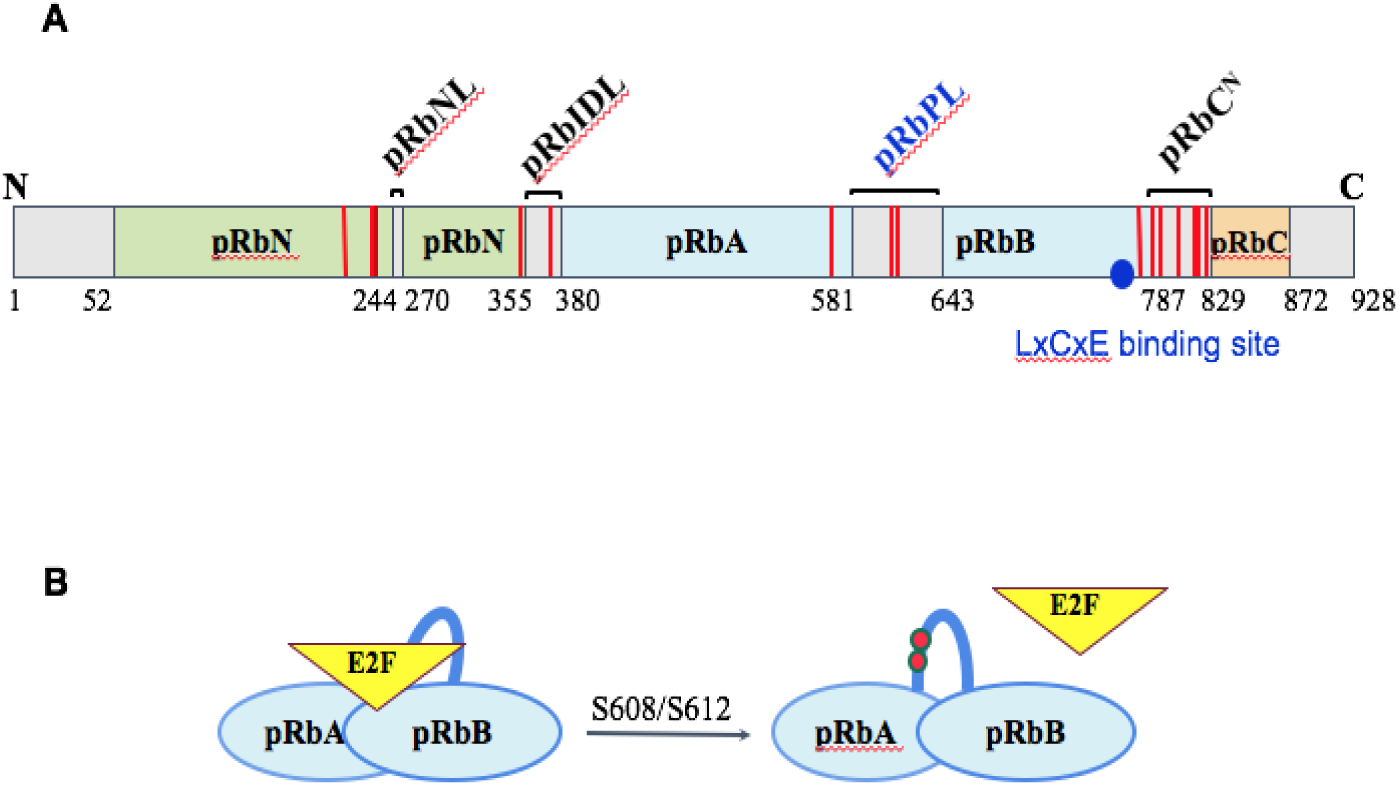
Domains and linkers of the retinoblastoma protein (pRb). (A) The schematic diagram representing the structured domains and intrinsically disordered regions between the structured domains. The phosphorylation sites are represented in red bars and the amino acid positions are numbered. The LxCxE motif binding site is represented as blue circle. (B) Model showing the impact of phosphorylation (red filled circles) at positions S608/S612 on the binding to E2F transcription factor transactivation domain to pRbAB domain.

pRb is a target of various post-translational modifications, the best understood of which is the phosphorylation. Thirteen of the sixteen potential serine-threonine phosphorylation sites identified in the protein are phosphorylated in cells, and all of these are present in the disordered linker regions [11]. Most of these sites are also conserved in human pRb isoforms, p130 and p107 indicating their functional importance (Fig. S1) [12]. Multisite, discreet phosphorylation events of pRb result in distinct domain rearrangements that inhibit its interactions with other protein partners [9].

A direct activation of pRb is an attractive strategy for controlling aberrant cell proliferation. Desired feature of any agent for achieving this will be to prevent the release of E2F by pRb even under phosphorylated conditions, and also upon binding with the LxCxE motif containing viral proteins. Feasibility of such an approach has been demonstrated *in vitro* [13]. However, the pRb construct used in this study lacked all flexible linkers, hence the effect of phosphorylation could not be assessed. One of the challenges for such approaches to be successful is to predict the *in vivo* behavior of the protein in response to the binding interactions, mutations, stability and function. Most of the pRb-AB crystal structures lack the RbPL linker between pRbA and pRbB. These structures show that the pRb-AB domain is rigid. *In vitro* studies of the pRb-AB domain along with RbPL linker show that the native structure has low *in vitro* thermodynamic and thermal stability that can be suppressed by its binding ligands, HPV E7 and E2F [14]. It may possibly have similar behavior *in vivo*. Additionally, other than a few low penetrance mutations, e.g., Arg661Trp and Cys712Arg [15, 16], most other mutants of pRb have not been evaluated in terms of their capacity for affecting local or large distance conformational changes that may have consequences for ligand binding and stability.

Molecular dynamic simulation is a powerful computational technique to study the conformational dynamics of a protein. An earlier study has reported MD simulation of pRb-AB without the connecting linker [17]. However, in order to build a credible model of pRb, structural information about its flexible linkers is needed. Protein linkers fulfill functions other than just connecting two domains of a protein, and can be instrumental for conformational and allosteric changes [18]. Accordingly, flexible linker regions and their alteration due to mutations can affect the allostery, stability and solubility of a protein [19, 20]. The linker between pRbA and pRbB, pRbPL has two phosphorylation sites, Ser608 and Ser612. A shortened pRbPL linker between pRbA and pRbB with phosphomimetic mutations corresponding to Ser608 and Ser612 has been reported earlier that shows a short helix formed by residues 602-607 [21]. Two residues in this helix align with the E2F side chains that make contact with the pocket domain. Based on this, and findings that the linker can compete with E2F in trans, it was proposed that phosphorylation of Ser608 and Ser612 directly inhibits E2F binding to the pocket domain pRbAB [21, 22]. Understanding the molecular mechanisms of ligand exchange at the E2F binding site of pRb is essential to delineate functional consequences of any mutation in the pocket domain and the linker region. In this context, it is noteworthy that of the ten hotspot missense mutations in pRb in cBioPortal database for cancer genomics (https://docs.cbioportal.org/), six are in the pRbAB and pRbPL linker. These six residues are Ser567, Tyr606, Leu607, Arg621, Arg661, Arg698 and Cys706.

The present work is an attempt to build a set of linker structures to help capture the conformational heterogeneity within the disordered pRbPL linker of pRb, and to understand the mechanism of Ser608 and Ser612 phosphorylation mediated release of E2F. In the absence of any structural information about the full linker, and hence, a suitable template, we have used *ab initio* modelling of the linker with the pRbA and pRbB boxes to develop a complete model of pRbAB with pRbPL linker. This model has been used to predict the mechanism of E2F release in response to phosphorylation of Ser608 and Ser612 in the pRbPL linker.

## 2. MATERIALS AND METHODS

### 2.1 Modelling and Evaluation of Retinoblastoma protein with linker

Retinoblastoma is a 928 amino-acid protein. The protein has N-terminal domains RbN-A and -B, and pocket domains pRbA and pRbB that are linked via several intrinsically disordered loops containing most of the phosphorylation sites in the protein (Fig. 1). To create a model of the pocket domain region pRbA and pRbB along with the 74 residues intrinsically disordered loop pRbPL linking these domains, the amino acid sequence of human pRb protein (Accession no. NP_000312) was retrieved in FASTA format from NCBI [23]. The pocket domain sequence covering 410 amino acids (380-790 aa) along with the pRbPL loop (581-640 aa i.e., 73aa) region was retrieved for developing an *ab-initio* model.

The two different tools, ROBETTA [24, 25] and I-TASSER [26]-[27] were used to build the models of pRbAB along with the pRbPL loop. ROBETTA primarily utilizes the homology modeling approach, followed by better template selection and modeling the missing loop region using the loop modeling algorithms to generate accurate 3D models of proteins. I-TASSER uses a combination of threading techniques that perform fold recognition with the database Protein Data Bank Similar to ROBETTA, it models the missing loop region that is subjected to selection of the best model based on a variety of metrics to evaluate the quality of the predicted structures, including root-mean-square deviation (RMSD), PROSA Z-score and Ramachandran plot criteria. Thus, both ROBETTA and I-TASSER rely on a combination of sequence-based information and structural templates to predict protein structures. The difference lies in the specific algorithms and strategies used for template selection, model building, and refinement of model protein. Both software have been extensively used successfully to develop ab-initio models of loops in proteins spanning a length of within 120 amino acids. Along with these two most common programs for ab-initio modeling, the predicted pRbAB structure from AlphaFold Protein Structure Database [28] was also considered [29,30]. The generated computational models were compared with retinoblastoma structures from the RCSB repository for evaluation of stereochemical quality and analysis of the interactions between the pocket domain residues. The PDB structures that were used for comparative analysis include an unliganded or apo-structure of pRbAB (PDB ID: 3POM), pRbAB bound to the transactivation (TA) domain peptide of E2F, referred hereafter as E2FTA (PDB ID: 1N4M) and a pRbAB structure with a shortened pRbPL linker containing the first 35 residues, i.e, aa 582-615 (PDB ID:4ELL) [6, 31],21]. Parameters derived from these structures are referred to as ‘reference dataset’ hereafter. The quality of models were evaluated using PROSA [32].

### 2.2 Molecular Dynamics simulation

MD simulations for the generated models of retinoblastoma structures were carried out using GROMACS 2019.4 with the AMBER99SB force field [33]. The simulation systems were prepared in cubic boxes filled with TIP3P water molecules. Neutralization was achieved with counter ions (Na^+^/Cl^−^), followed by energy minimization using the steepest descent method. The systems underwent 100 ps NVT and NPT equilibration steps before a 100 ns MD simulation, saving energy data every 10 ps.

### 2.3 Phosphorylation of Rb protein

Phosphorylation was introduced using the PyTMs plugin [34, 35], focusing on residues 608 and 612 in the pRbPL linker connecting the pocket domain. A targeted approach was employed by modifying the “Selection” parameter from “all” to “sele”, ensuring exclusive phosphorylation of the selected residues. Subsequently, molecular dynamics (MD) simulations were conducted to investigate conformational dynamics. Notably, phosphoserine parameters were incorporated into the amber force field file. We prepared the system with phosphorylations at Ser608 and Ser612. These modifications were introduced manually based on the known phosphorylation sites.

### 2.4 Mutation

To investigate the structural and functional consequences of point mutations, the modeled protein structure was subjected to site-directed mutagenesis at arginine 661 (R661), replacing it with tryptophan (W) to generate the R661W mutant. This mutation was introduced using the mutagenesis tool available in the PyMOL Molecular Graphics System [36]. The most probable rotamer for tryptophan was selected based on minimal steric clashes and optimal geometry. Following the mutation, the structure was energy-minimized to relieve any unfavorable contacts introduced during the mutation process.

### 2.5 Docking with E2FTA domain

To evaluate the interactions between the generated models and E2F, molecular docking was carried out with E2FTA domain peptide (410-427 aa) to validate their consistency with reported interactions. Ligplot+ was used to generate 2D interaction maps [37]. Additionally, the impact of phosphorylation at Ser608 and Ser612 on interaction with the E2FTA was studied to comprehensively understand the mechanism of interaction.

The binding pocket was described by the interacting residues reported in PDB: 1N4M and from literature [17]. Docking was carried out by utilizing the High Ambiguity Driven protein-protein docking server (HADDOCK ) [38].

## 3. RESULTS

### 3.1 Quality assessment of computational models of pRbAB with pRbPL loop

The quality of five models from both I-TASSER and ROBETTA, as well as the AlphaFold model were analyzed in order to determine the most accurately modeled structure(s). The models generated by I-TASSER exhibited low C-score values ranging from -3.29 to -0.80, indicating a lack of confidence in their predictions. On the other hand, the ROBETTA models scored a confidence level of 73 on a scale of 0 to 100, suggesting higher quality in the generated models. Among the ROBETTA models, the first two demonstrated the least error approximation compared to the remaining three. In case of the AlphaFold model, the confidence scores for the loop region were below 50 on a scale of 0 to 100, while the pocket region displayed confidence levels exceeding 90%. These scores, specific to each modeling software, reflect how well the predicted structures align with both multiple sequence alignment data and known PDB structure information, serving as a fundamental indicator of model accuracy (Table S1).

All generated models were further evaluated based on structural examination of the models to make sure the generated models are devoid of steric clashes. The I-TASSER models showed almost 25% residues in the unfavorable regions of the Ramachandran plot suggesting poor torsional angles and steric clashes (Table S1). In contrast, the Robetta and AlphaFold models showed most of the residues in the favorable region of the Ramachandran plot (Table S1). The spatial arrangement and interactions with neighboring residues were evaluated by Verify3D for each model by comparing the energy score of each residue in the modeled structures to the distribution of scores for that type of residue in the reference dataset. Most residues (>80%) had scores within 0.2 when the complete pocket domain with the linker was considered in models generated by ROBETTA and AlphaFold, while the I-TASSER models showed more fluctuations in scores in the loop region.

The generated models were also analyzed using PROSA to check whether the *z*-scores of the generated structures are within the range of scores typically found for native proteins by comparing with the z-scores of the pRbAB structures 1N4M (without pRbPL loop) and 4ELL (with shortened pRbPL loop). The z-score of 1N4M, is -6.97Å. The z-scores of the models generated by Robetta were -6.97+/- 0.5, while the z-scores of the I-TASSER models were in the range of -6.97+/-1.6. The Z-scores of the models compared to 4ELL, that has a shortened loop, showed a more negative value of z-scores compared to those obtained with all I-TASSER models (Table S1) indicating better quality of the models generated by Robetta and AlphaFold compared to those generated with I-TASSER.

The fragment-wise RMSDs were calculated and compared with crystal structures. The resolution of 3POM is 2.50Å, while that of 1N4M is 2.20 Å [31],[6]. Thus, both structures are of good and comparable resolution. I-TASSER models showed a reasonable range of RMSD values varying from 1.1+/-0.3Å against 3POM and 0.90+/-0.60 Å against 1N4M (Table S1). Robetta models showed a RMSD value of 1+/-0.1 Å against 3POM and 0.9+/-0.1 Å against 1N4M structure. The AlphaFold model showed even lower RMSD values with respect to both the crystal structures, surpassing the other two models in accuracy.

Based on these analyses, ROBETTA models 1 and 2 and the AlphaFold model (referred hereafter as Model3), were taken for further analysis. The RMSD values of the models generated by Robetta and AlphaFold were also compared with a previously published pRbAB structure with a short linker (PDB ID:4ELL) that has a resolution of 1.98Å [21]. RMSD values were compared with the linker region of the models that showed values of 0.783, 1.049 and 0.889Å for Robetta models 1 and 2, and AlphaFold model respectively, suggesting only a low deviation from the published structure. Models generated by Robetta and AlphaFold have a higher proportion of residues residing within the permitted region of the Ramachandran plot, accompanied by low z-scores relative to 4ELL. This suggests the absence of clashes in the models and indicates comparability with crystal structures of similar dimensions. Conversely, although I-TASSER models demonstrate favorable RMSD agreement with 3POM and 1N4M structures, they exhibit markedly higher z-scores compared to 4ELL, indicative of diminished similarity to crystal structures. Moreover, the I-TASSER models exhibit low Ramachandran favorability, with numerous residues occupying unfavorable regions, hinting at potential clashes and reflecting poorer model quality, thus justifying their exclusion from further consideration.

Comparison of the 3D structures of these models with the selected pRbAB structures is shown in Fig. 2A. The 4ELL structure that has a shortened pRbPL loop and a phosphomimetic mutation of Ser608 shows a helix formed by residues 602-607 that is similar to that formed by the C-terminal helix of E2F [21]. This helix was observed in all the three selected models, even without any phosphomimetic mutation of Ser608. All three models showed the linker fluctuating between the short helix or a loop (Fig. 2B) showing that the linker has a propensity to form this short helix independent of the phosphorylation status of Ser608.

**Fig. 2:**
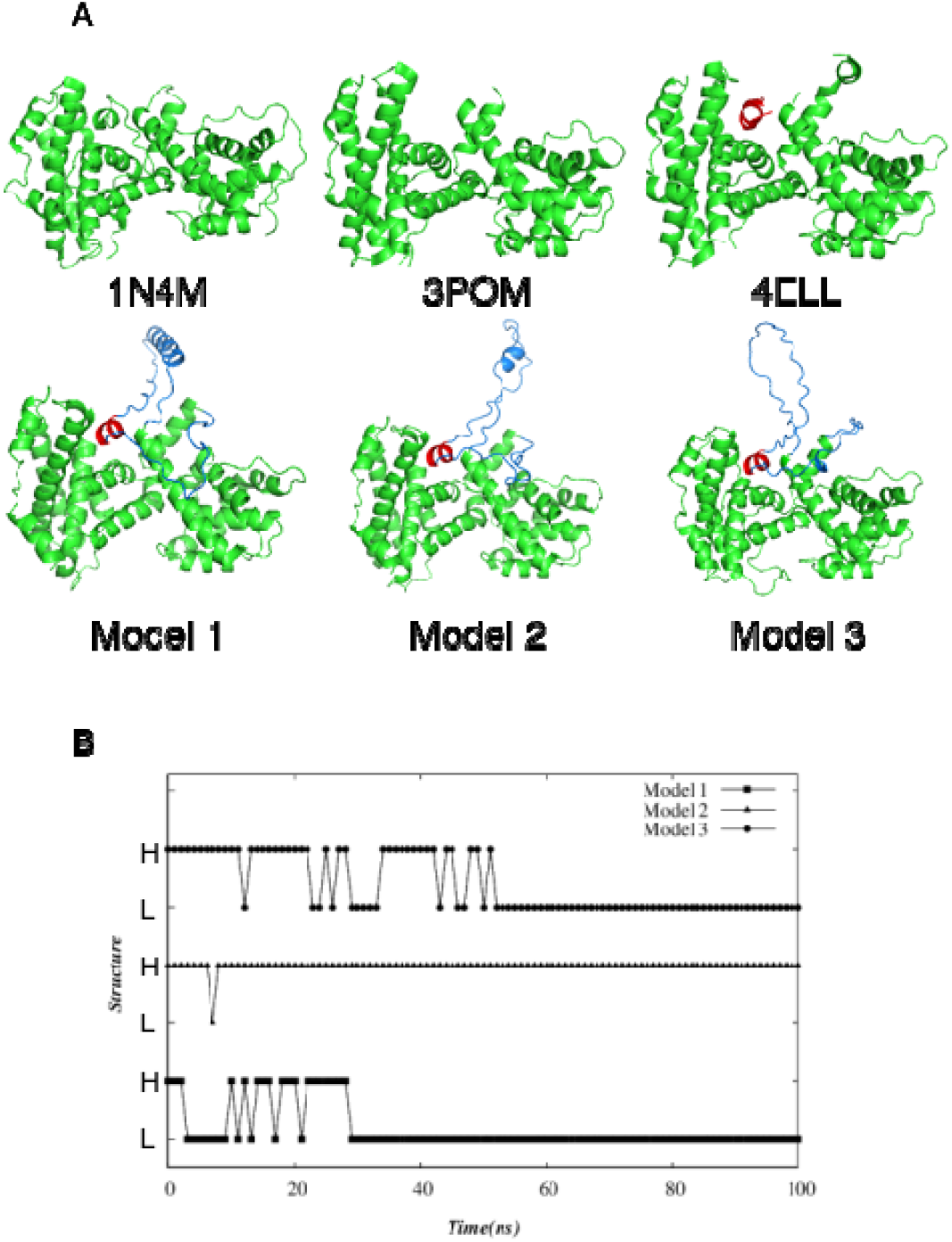
The 3D visualization of generated models. (A) PDB structures used for comparison 1N4M, 4ELL and 3POM are shown for reference. The short helix present in 4ELL structure is shown in red. The selected models with complete loop generated by ROBETTA (Models 1 and 2) and from AlphaFold (Model 3) showing the pRbPL loop (blue) and the short helix present in all three models (red). (B) The graph shows fluctuation of the linker with or without the short helix in a 100 ns simulation of the linker in each model. H, helix; L, loop.

To assess whether the loop has any significant effect on the molecular interactions between pocket A (380-580) and pocket B (641-790), the interactions in the generated models were compared to the PDB structure of unliganded pRb-AB (PDB: 3POM), pRb-AB bound to the E2FTA (PDB: 1N4M), and pRb-AB with partial loop with phosphomimetic mutation at S608 (PDB:4ELL). All the three structures show conservation of most of the interactions between the pocket domains (Fig. S2). Thus, interactions between Glu559 with Arg661 and His699; Ser565 and Tyr563 with Asp697; and several non-bonding interactions of Phe650 with pocketA residues are conserved in all three PDB structures (Fig. S2). However, the interactions are not identical in the three structures as expected from slight changes depending upon the binding partners and presence or absence of part of the pRbPL linker. Thus, presence of even a shortened linker and introduction of a single phosphomimetic mutation results in subtle changes in interactions between the pocket domains without substantial conformational changes. Comparison of the interactions between the pockets in the generated models with full length pRbPL linker showed conservation of all interactions mentioned above including a majority of the salt-bridge and non-bonded interactions. Thus, the presence of the linker did not cause any major perturbations in the interactions of the A-B pockets in the generated models (Fig 3A). However, similar to the differences between 1N4M and 4ELL, the three models showed some differences between themselves and with the model structures, as can be expected given the flexibility introduced due to the long loop. Next, the interactions between the pRbPL loop and the pocket domains was examined (Fig 3B). It has been repo ted that the critical interactions within the partial loop structure, specifically the salt bridge between Asp604 and Arg467, and the van der Waals interaction between Tyr606 and Ile481, and Phe482 that overlap with interactions made by the C-terminal E2F helix form the basis for a competitive mechanism for E2F release upon phosphorylation of Ser608 [21]. While unphosphorylated models 1 and 2 did not show Asp604 participating in any interactions with the pocket domains, unphosphorylated model 3 showed a H-bond between Asp604 and Arg467 (Fig. S2). The Tyr606 consistently showed non-bonded interactions with Phe482 in all the three models, but interaction with Ile481 was not observed in model 1. Thus, the pRbPL linker contacts residues in both pockets even without phosphorylation but does not perturb the pocket AB interactions to any significant extent, ruling out large-scale movement of pocket domains with respect to each other.

**Fig. 3:**
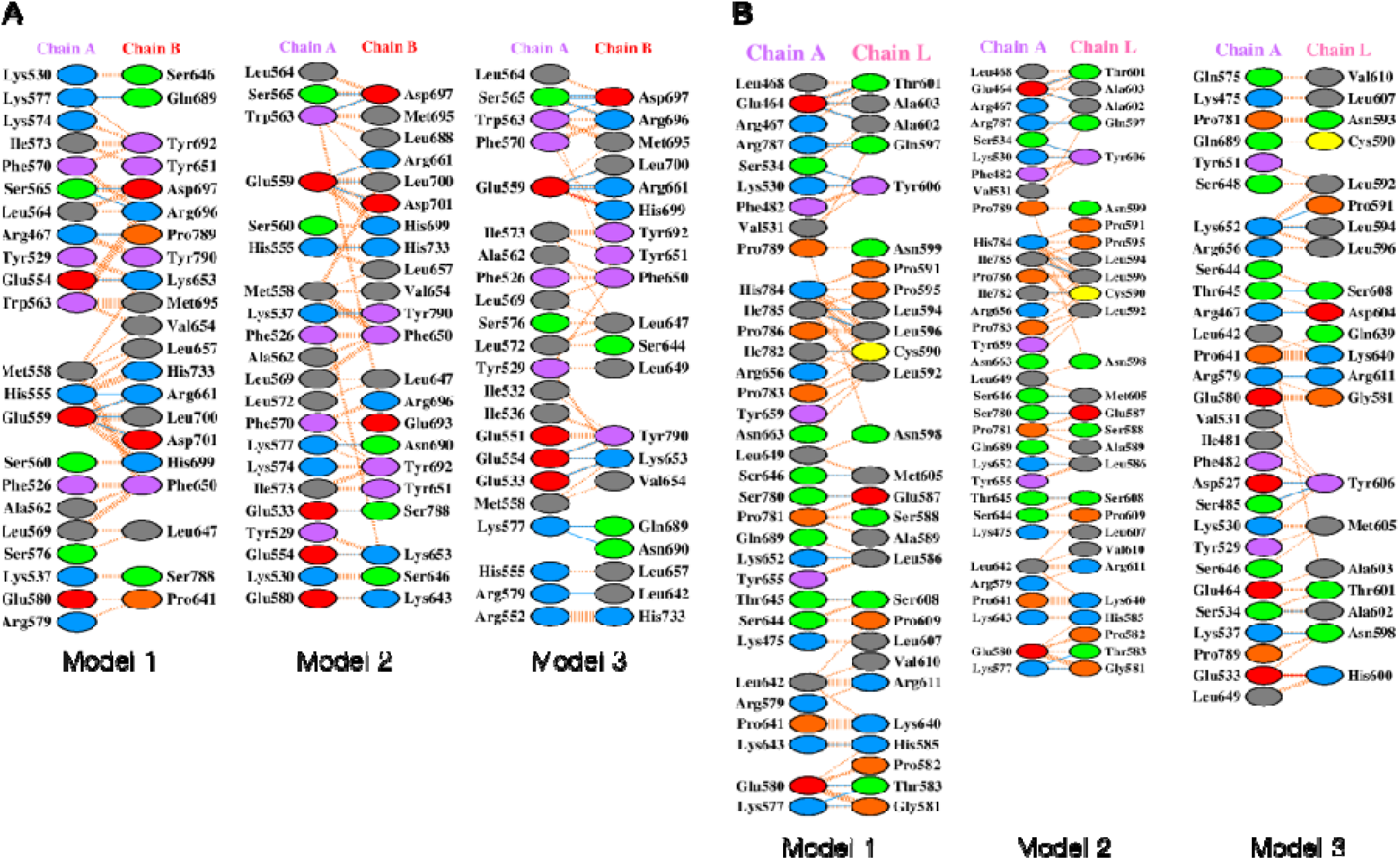
Ligplots of the interacting residues. (A) Between pocket A (Chain A) and B (Chain B) in the modelled structures. (B) Interacting residues between pRbAB pocket domains A, B (Chain A) with the pRbPL linker (Chain B). Color code for the amino acids is Grey: Neutral; Blue: positively charged; Red: negatively harged. Interactions via the side chains are shown in Green: neutral side chain interactions; Violet: hydrophobic; Yellow: Special cases (Gly,Pro,Cys). Nonbonded interactions are shown with dashed orange and sky blue lines and bonded interactions are shown with continuous red lines).

### 3.2 Effect of pRbPL loop phosphorylation on the dynamics of the pocket domains

In order to analyze the conformational changes in the three models, Molecular Dynamics (MD) simulation studies were performed over 100 ns simulation time using Gromacs. The Root Mean Square Deviation of protein backbone (RMSD) of Model 1 fluctuated between 0.15-0.51 nm showing a more stable and compact structure with a radius of gyration (Rg) between 2.35-2.48 nm compared to the other two models that showed much higher fluctuations: Model 2 showing 0.17-0.72 nm and Model 3 showing 0.16-0.83 nm RMSD, with Rg values being 2.32-2.54 nm for Model2 and 2.41-2.57 nm for Model 3 (Fig. 4A, B). The RMSD of the first two models were more stable after 80 ns and the Rg of all the three models were stable after 80 ns.

**Fig. 4:**
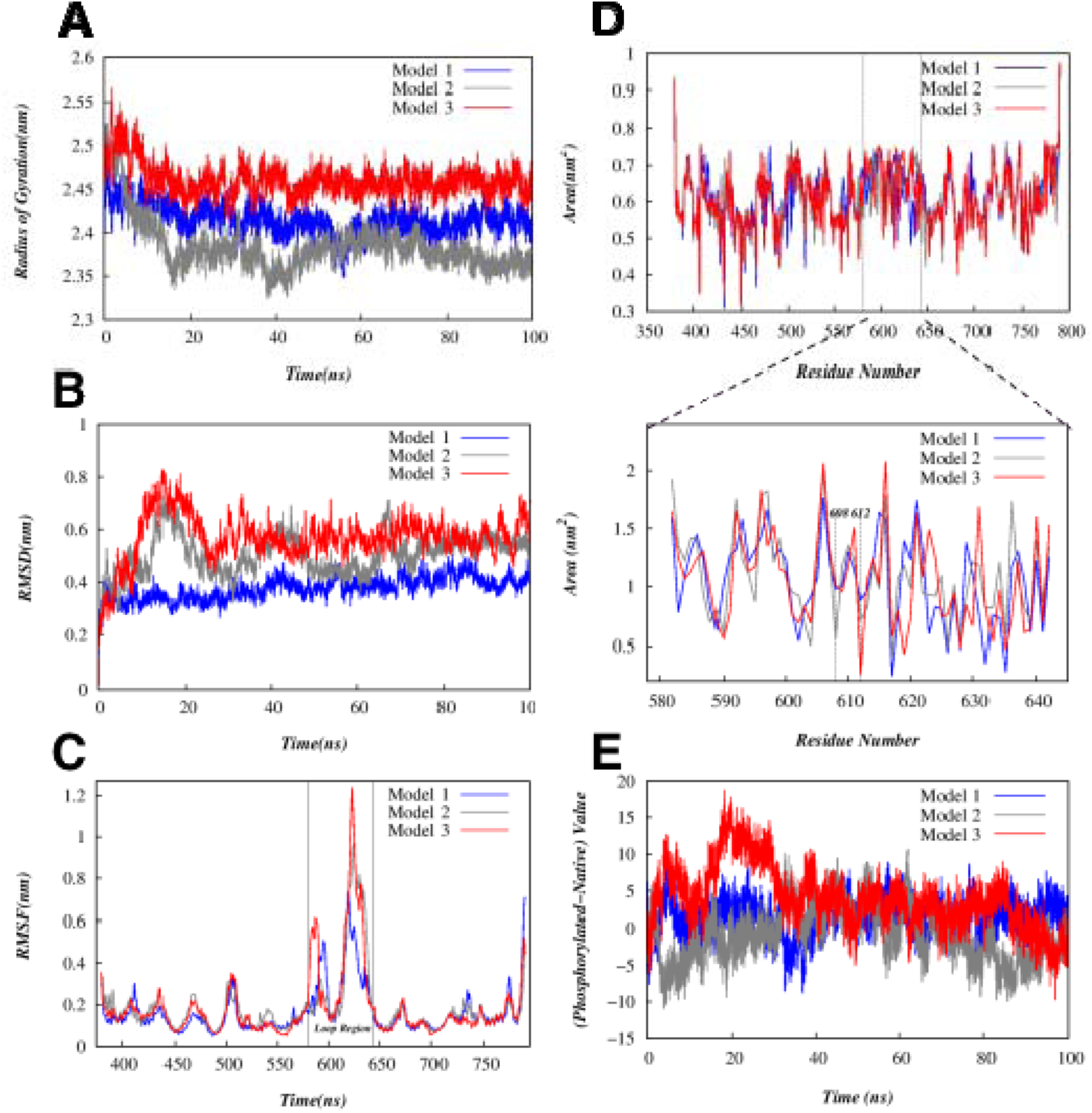
Dynamics of the pRbPL loop region. (A) The Root Mean Square Deviation (RMSD) along the backbone of the unphosphorylated models. (B) the Radius of Gyration of the unphosphorylated models. Root Mean Square Fluctuations (RMSF) and Solvent Accessibility (SASA) of unphosphorylated models. (C) RMSF per residue of the protein throughout the simulation time. Vertical lines mark the residues within the pRbPL loop. (D) SASA of the unphosphorylated models throughout the simulation. Expanded region, aa 580-650 of the pRbPL loop throughout the simulation time is shown below. Serine residues at 608 and 612 positions are marked. (E) Time dependent variation in the difference in the surface accessibility of the phosphorylated and native protein within the loop region.

The Root Mean Square Fluctuation per residue showed the pocket domain fluctuations to be low and comparable in all three models (Fig. 4C) agreeing with previous reports that the pocket domains are rigid structures [31]. In contrast, the RbPL loop region in all three models had considerably higher fluctuations compared to the pocket domain region, with models 2 and 3 showing much higher fluctuations than model 1 (Fig. 4C). This is in keeping with the disorder and flexibility expected from a loop. Significantly, within the loop, the region containing Serine608 and 612 had the highest mobility (Fig. 4C). It is expected that due to its flexibility and dynamics, the RbPL loop will also be accessible to the solvent. This was confirmed by measuring the solvent accessibility areas that were observed to be elevated near the loop region (Fig. 4D). These features may allow for a higher probability of undergoing phosphorylation due to these fluctuations that can briefly expose these residues to the responsible kinases.

Phosphorylation at positions 608 and 612 has been reported to compete with E2FTA binding by interaction of the loop residues with the pocket domain residues [21, 22]. The 4ELL structure that had a shortened linker with only twenty-four linker residues, and had phosphomimetic mutations of Ser608 and Ser612, was shown to have a short helix formed by the residues 602-607, similar to that formed by the C-terminal of E2F and making the same contacts with pocket residues as by the E2F helix [21]. The phophomimetic Ser608Glu was found to interact with the helix aP11 by H-bonding with amide protons of Ser644 and Thr645, and side chain hydroxyls of Ser644 and Ser646 [21]. While interactions between unphosphorylated Ser608 and Ser644, Thr645 were observed in all the three models, phosphorylation at Ser608 and Ser612 followed by MD-simulation for 100 ns showed increased accessibility of the loop region compared to unphosphorylated Ser608/Ser612 (Fig. 4E). Therefore, in the context of the full linker, phosphorylation of Ser608 and Ser612 results in more solvent exposure and these residues may not be buried in the pocket groove.

### 3.3 Model Validation and Structural Analysis: Conformational patterns in the E2F Binding interactions

Although experiments using the RbPL loop in trans have suggested competitive mechanism for release of E2F by phosphorylation of Ser608 and Ser612 [21, 22], the flexibility and conformational dynamics of a linker whose ends are tethered to the protein may pose some constraints to the volume that is available for the transactivation domain to approach and bind to the cleft. Therefore, accessible interfacial area and volume of the cavity between the AB-pockets were examined using CastP software [39]. The surface area of the interfacial region between the A and B pockets, as well as the volume of the cavity formed within these pockets across various cluster structures obtained from each model was measured (Fig. 5) [40]. The volume of the E2FTA is measured at 1745.512 A^3, excluding any void volume. Model 2 exhibits ample space and volume between pockets A and B, encompassing the loop portion. In contrast, both Model 1 and Model 3 demonstrate a less accessible surface area between pockets A and B, making it challenging to accommodate the E2FTA (Fig. 6). Consequently, further investigations were conducted with Model 2 to elucidate the fluctuations in loop conformation, in order to understand the molecular mechanisms involved in E2F binding to the pRb pocket domain.

**Fig. 5:**
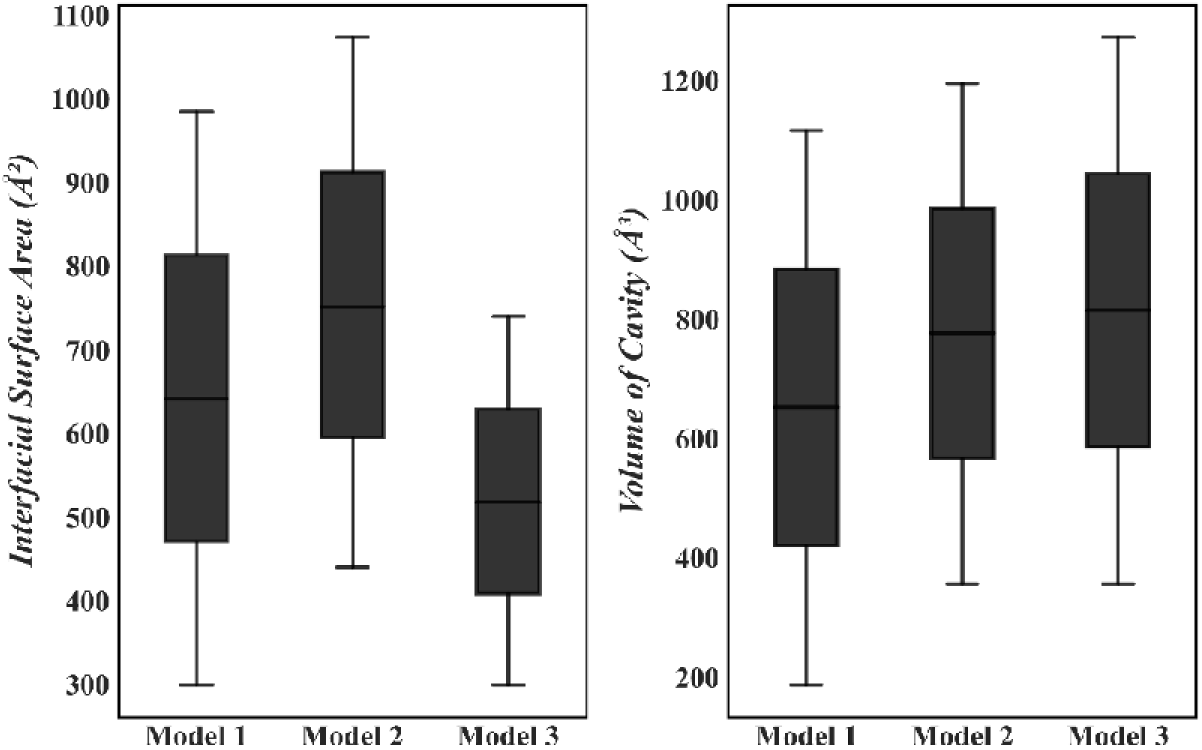
Estimation of the interfacial area and volume of the pocket cavity. The box plots illustrate the distribution of interfacial surface areas (left) and volumes of cavity (right) for the three models. Each boxplot visually encapsulates the range of interfacial surface areas and volume of the cavity between the pockets, with the central line depicting the median value. Whiskers extending from the boxes denote the range of the data, excluding outliers.

**Fig. 6:**
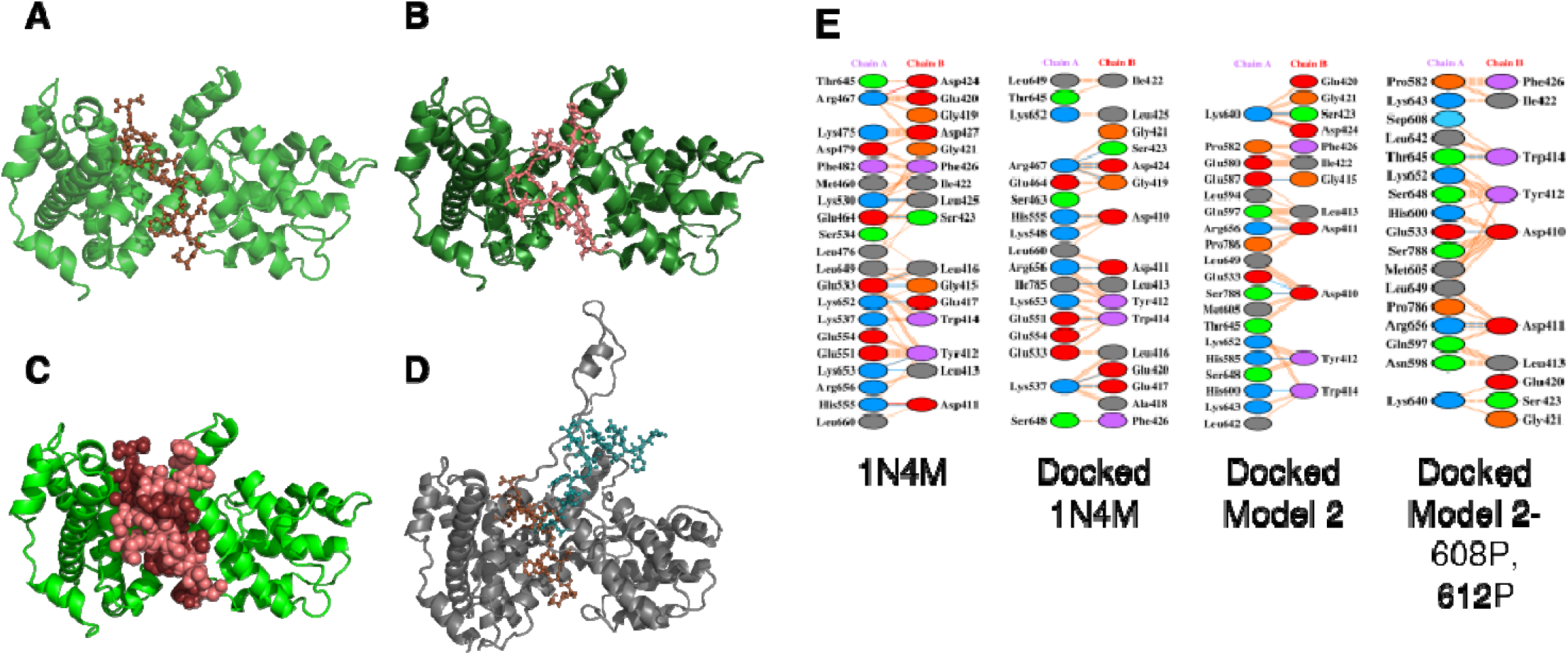
Interaction of E2FTA with the AB pocket of pRb. (A) The crystal structure 1N4M depicting E2F TA interactions with the pocket domain. E2F peptide is shown in purple with 1N4M in green. (B) The docked E2F peptide on 1N4M structure using HADDOCK. The E2FTA is shown as stick representation in salmon. (C) Alignment of 1N4M crystal and docked structures. (D) E2FTA (Teal) docked to Model 2. For comparison, E2FTA in 1N4M structure is shown in purple. (E) The LigPlot+ analysis of E2FTA interactions at the AB interface in 1N4M crystal structure, HADDOCK docking of E2FTA to 1N4M, unphosphorylated and phosphorylated Model 2. Chain A, pRb; Chain B, E2FTA. Color code for the amino acids is Grey, Neutral; Blue, positively charged; Red, negatively charged. Interactions via the side chains are shown in Green, neutral side chain interactions; Violet, hydrophobic; Yellow, Special cases (Gly,Pro,Cys). Nonbonded interactions, dashed orange lines; H-bonds, blue solid lines; salt-bridge, solid red lines.

The interacting residues in pRb pockets A and B with E2FTA have been described in the crystal structure, PDB ID:1N4M. Using the HADDOCK tool, we examined the interactions between pRbAB and E2F residues. In comparison to the crystal structure, the docked complex showed differences in the placement of N-terminal of E2FTA that is oriented more towards the pRb-pocket B as opposed to the A-pocket in the crystal struct re (Fig. 6A, B, C). A slight reduction in the number of hydrogen bonds was also observed in the docked structure, with 10 hydrogen bonds identified in the docked structure versus 13 in the original crystal structure (Table 1, Fig. S2).

**Table 1:**
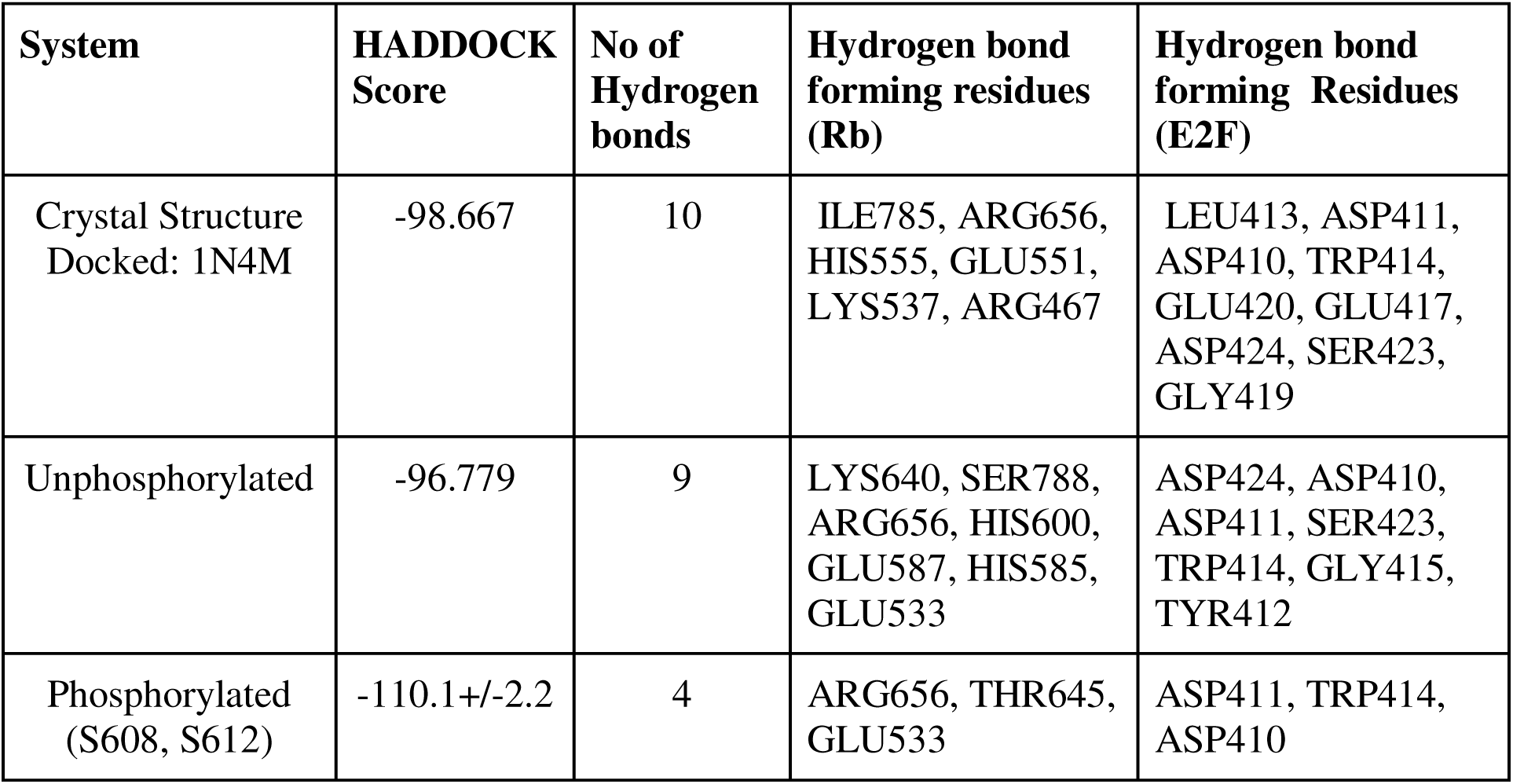
Docking scores of the Model2 with the E2FTA under different phosphorylation states.

Nevertheless, several interactions were found to be common in both the crystal and docked structures (Fig. 6E). For example, Arg467 of pRbAB forms hydrogen bonds with Asp424 and Gly419 of E2FTA in both structures, showing similar geometry. Other pRb residues were also found to be common between the docked and crystal structures involved in bonded and non-bonded contacts. Thus, Lys537, Lys652, Leu649, Glu554, Glu551, Leu653, Arg656, and Ile785 were involved in interactions with E2F in both the docked and crystal structures.

Although the specific E2F residues may differ slightly in each case, these pRbAB residues play a role in maintaining the binding interface. Since Model 2 exhibited ample spacing between pocket A and B Fig. 5), potentially facilitating E2F binding compared to the other two models, and did not display any atypical changes in the dynamics analyses (RMSD, Rg, etc.), it was selected for further investigation.

Docked structures are not likely to always capture the experimentally determined pose, and differences are expected. However, preservation of most of the interacting residues and interactions shows that the docked structure recapitulates the observed interactions observed in the crystal structure and can be used to examine the docking of the E2FTA to modeled structures. This docking strategy ensures that critical interaction points are maintained in the docking process. As opposed to 1N4M structure where the E2FTA is bound snugly be ween the A and B pockets of pRbAB, in the modeled structure with the loop, the penetration between the cleft is restricted at the edges of the regions close to the loop (Fig 6D). A number of interactions between the E2FTA and the loop residues, in the regions between 570–610 and 635–645 were observed in the docked structure (Fig. 6E). The complex formed nine hydrogen bonds, comparatively fewer than in the crystal structure, but with a similar number of non-bonded interactions (Table 1).

Nevertheless, several common interactions were observed between the crystal structure and Model 2 (Fig. 6E). For instance, Asp411of E2FTA interacts with pRbAB Lys548 in 1N4M, and with Arg656 in Model 2, indicating that the same Asp residue is involved but with different interacting partners. Similarly, Glu533 of pRb interacts with Leu416 and Gly415 of E2FTA in the crystal structure, while it interacts with Asp410 in Model 2. Given the dynamic nature of the loop, it is expected that E2FTA will not be able to penetrate deep into the interdomain space as in the crystal structures that do not contain the loop. However, almost similar number of interactions between E2FTA and Model 2 to that observed in 1N4M suggest that the binding affinity is not drastically reduced in the presence of unphosphorylated pRbPL loop.

### 3.4 Effect of phosphorylation of pRbPL loop on conformational flexibility and E2F interaction

The E2F was docked to model2 using the unphosphorylated binding pocket as a reference. In the unphosphorylated pRbAB-E2FTA complex, pRbAb Lys640 makes a salt bridge, a H-bond and several non-bonded interactions with E2FTA. In contrast, in the phosphorylated protein the salt-bridge and H-bond of Lys640 are not observed. The most consistent interaction between both states was the bond between Arg656 of pRbAB and Asp411 of E2FTA, highlighting it as a crucial interaction point. The reduction in hydrogen bonds suggests weaker or less frequent interactions of E2FTA with the pocket domain upon phosphorylation at Ser608 and 612. Though the direct impact of these residues with E2F was not observed. Correspondingly, the HADDOCK score was further increased upon phosphorylation of both Ser608 and Ser612 (Table 1), thus agreeing with the published results that phosphorylation of these residues leads to E2F release.

To understand how phosphorylation affects the conformational dynamics and charge distribution within the critical RbPL loop region, we examined the end-to-end distance of the loop region and assessed its electrostatic properties. The unphosphorylated pRbAB models do not show much fluctuation and change in the end-to-end distance throughout the simulation time of 100 ps. In contrast, phosphorylation of both Ser608 and Ser612 in all the three models exhibited higher fluctuations and an overall increase in the end-to-end distance in the loop region, indicating potential structural rearrangement (Fig. 7A). Further, determination of the electrostatic potential as a function of distance from phosphorylated Ser608 and Ser612 showed changes to a higher positive value compared to native ones, suggesting altered charge distribution along the pRbPL upon phosphorylation (Fig. 7B).

**Fig. 7:**
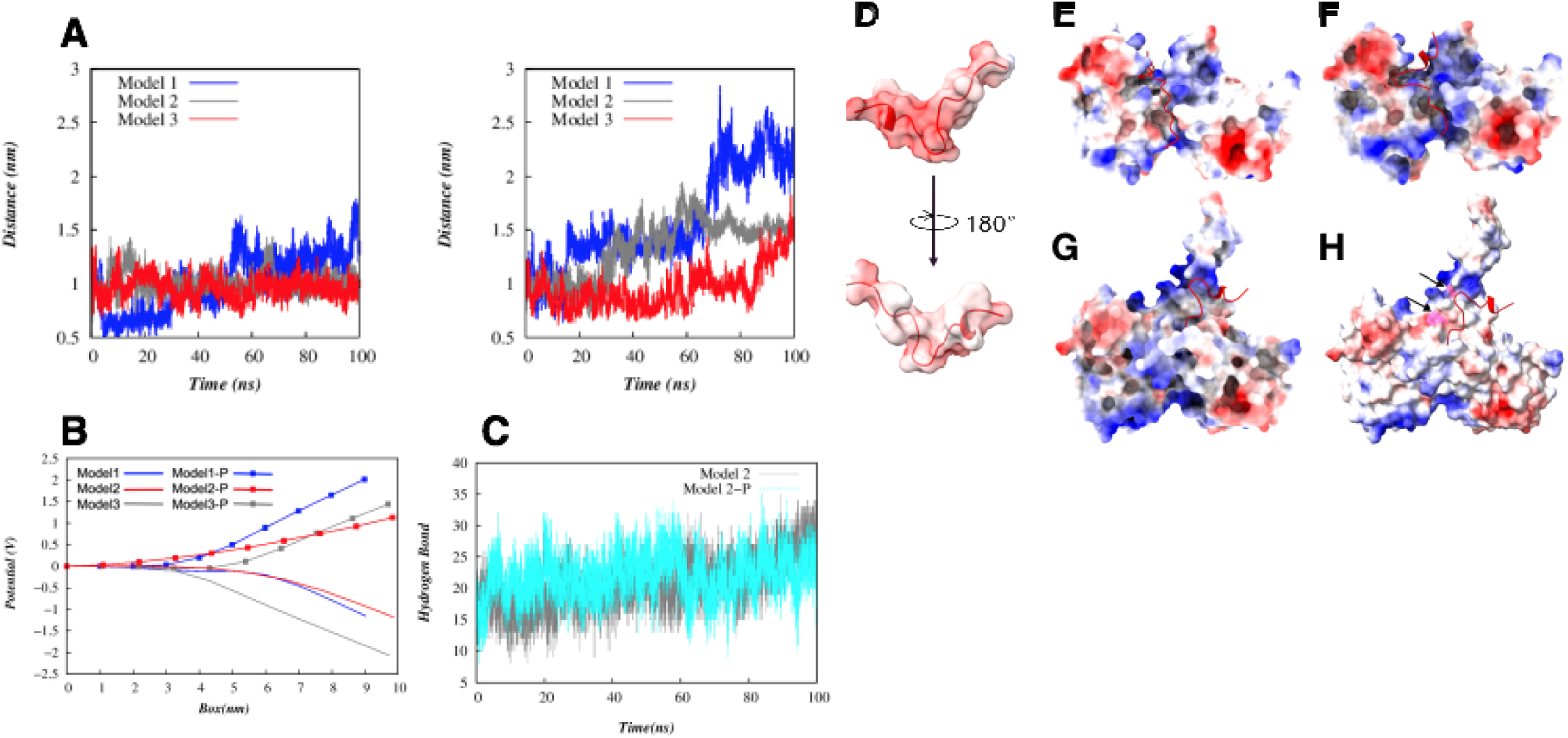
RbPL loop end-to-end distance fluctuations and electrostatic potential of phosphorylated and non-phosphorylated pRbAB models. (A) Comparison of fluctuation of loop end-to-end distance of the non-phosphorylated (left) and phosphorylated (right) pRbAB models. (B) Electrostatic potential trends of RbPL loop across the box under different phosphorylation states. (C) MD simulation of the number of Hydrogen bonds formed in phosphorylated (Cyan) and non-phosphorylated (Grey) pRbAB model 2. (D) Space-filling structure of E2F TA peptide showing the electrostatic property. Backbone structure. Electrostatics of 1N4M (E) crystal structure with E2FTA peptide (Red cartoon) and (F) HADDOCK docking. Electrostatics of (G) Unphosphorylated Model 2 with the linker and (H) S608, S612 Phosphorylated model2 with the linker docked to E2FTA peptide. The E2FTA peptide is shown in red color as cartoon. Pink, phosphorylation sites, indicated by black arrows.

Phosphorylation can change the local and global electrostatic properties of a protein that can impact the conformational behavior of a protein with the effects depending on the specific protein [41]. Our findings suggest that phosphorylation of the RbPL loop results in a more dynamic movement of the loop and an increase in the electrostatic potential of the loop. Further analysis of hydrogen bonds formed during the simulation revealed a decrease in the number of H-bonds in the phosphorylated structures (Fig. 7C), suggesting changes in the structural stability and dynamics, that can impact pocket domain-pRbPL linker interactions. This underscores the need for additional investigation to elucidate the specific functional consequences of these changes.

We further validated our modeled structure by introduction of the highly frequent mutation in pRb, R661W. This mutant which is temperature sensitive, partially inactivates pRb function and has significantly reduced binding with E2F1 [42]. Introduction of this mutation resulted in a reduction of surface accessibility, particularly within the pocket cavity, with values ranging around 421.451 +/- 200. Simulation of the mutated protein for 100ns showed a slight increase in the Root Mean Square Fluctuation (RMSF) of Cα analysis in the mutated R661W structure within the loop regions compared to the native one, particularly in proximity to loop regions (Fig. 8A). The time-dependent analysis of the whole protein Solvent Accessible Surface Area (SASA) reveals an overall decrease in solvent accessibility of mutated model over the entire simulation period (Fig. 8B). Additionally, when examining SASA changes on a per-residue basis, noticeable fluctuations are observed. The solvent accessibility of the loop region is decreased at ends in case of R661W when compared to the wild type (Fig. 8D).

**Fig. 8:**
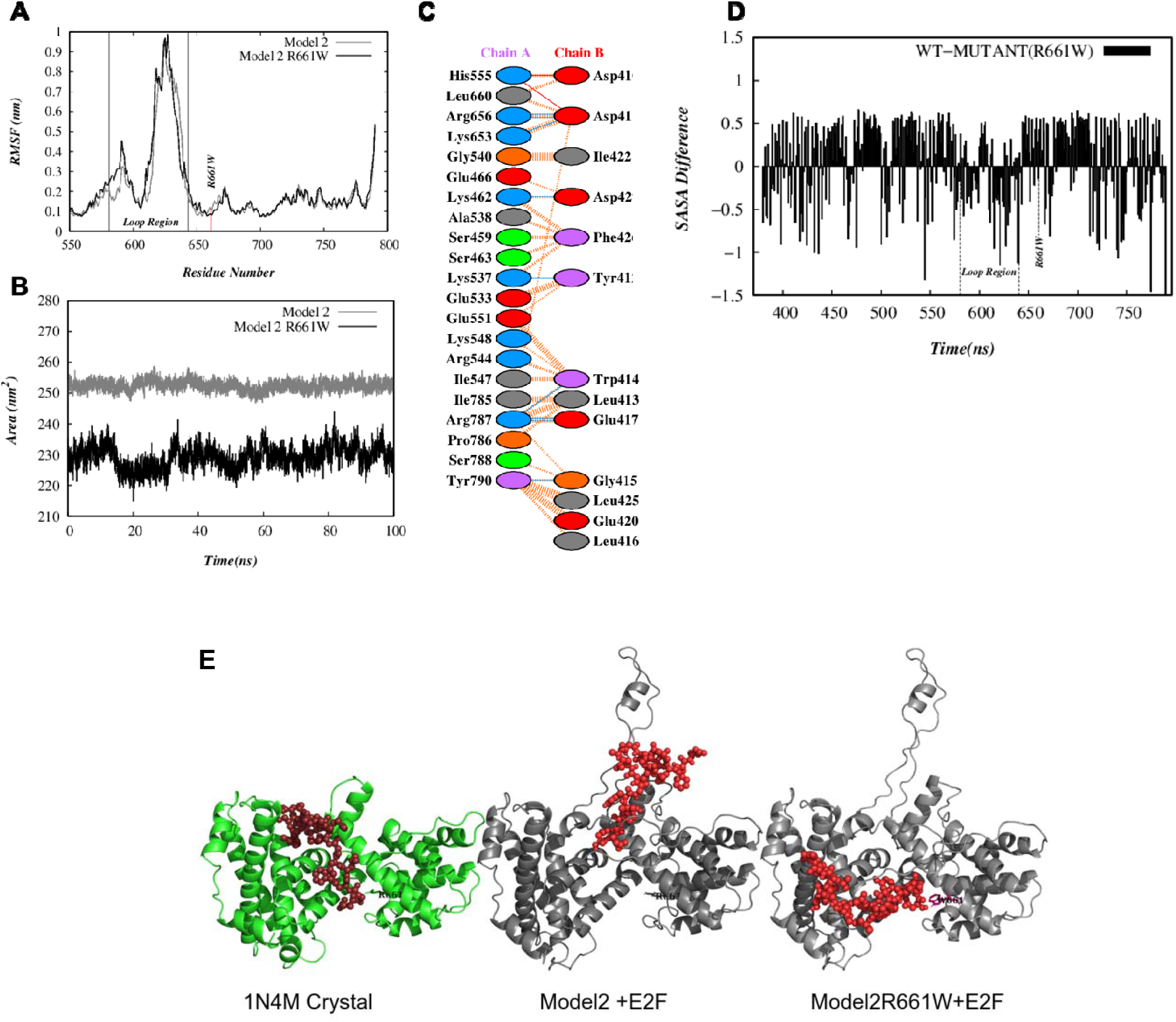
Changes in SASA and interactions with E2FTA in R661W mutant. **(A)** Root Mean Fluctuation and (B) Solvent Accessibility of R661W mutant. (C) Ligplot showing interactions between R661W pocket domain (ChainA) with E2FTA (Chain B). Nonbonded interactions, dashed orange lines; H-bonds, sky blue lines; bonded interactions, continuous red lines. (D) Percentage change in Solvent Accessible Surface Area (SASA) of the R661W model in comparison with the native counterpart. (E) Docking of E2FTA to 1N4M, Model2 and R661W mutant. The mutated position is indicated with an arrow.

Upon docking of the E2FTA peptide to R661W, the N-terminal region of E2F exhibited interactions similar to those observed in the crystal structure, albeit with a different orientation from both the crystal structure and the full-length model2–E2FTA complex (Fig 8E). However, the C-terminal region of E2FTA formed additional interactions with the pocket A domain of Rb. In the 1N4M crystal structure, His555 of pRbA pocket forms a salt-bridge and a H-bond with Asp411 of E2F, while in the docked 1N4M, this interaction is via a H-bond and a non-bonded interaction (Fig 6E). Both full length model 2 and phosphorylated model 2 do not show His55 making any contact with E2FTA, possibly due to the loop (Fig. 6E). In the R661W-E2F complex, His555 interacts with both Asp410 and Asp41 via salt-bridges (Fig. 8C). However, in the mutated complex, the E2FTA does not bind to the cavity between the pocket domains and shows more surface interactions (Fig. 8E). Additionally, a comparison of the docking scores shows that although the number of H-bonds do not change between the unmutated model 2 and R661W mutant docked to E2FTA, the binding energy of the mutant was -94+ -8.7 kCal/mol (Table 2) as opposed to -101.2+-2.2 kCal/mol for unphosphorylated model2 and -82.0+ -3.0 kCal/mol for phosphorylated model2 (Table 1). These results show that while the phosphorylated model 2 has a higher binding energy than unphosphorylated model 2, the R661W may have a more fluctuating binding energy landscape.

**Table 2:**
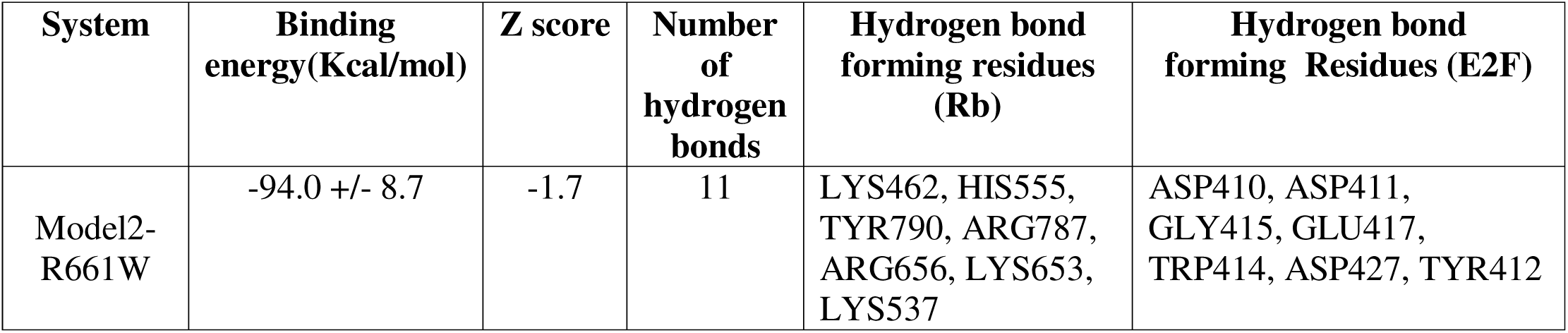
Docking scores of the Model1 R661W mutant with the transactivation domain of E2F.

To further understand how the R661W may have a lowered affinity to E2F, we examined the relationship between the SASA of the loop compaction of the protein and the surface free energy. While unphosphorylated model2 showed a relatively compact and structured free energy landscape, with one dominant low-energy region and a secondary, less populated state, the R661W mutant showed a more expanded energy landscape, with a broader distribution of states, indicating increased flexibility or multiple conformations (Fig. 9). Phosphorylated model 2, on the other hand showed a strikingly different distribution that appears to be bimodal, with two distinct low-energy basins, suggesting that phosphorylation induces significant conformational shifts (Fig. 9). These results show that the R661W mutation leads to a more dispersed conformational ensemble. While phosphorylation introduces two major low-energy states, implying possible conformational switching between different functional forms.

**Fig. 9:**
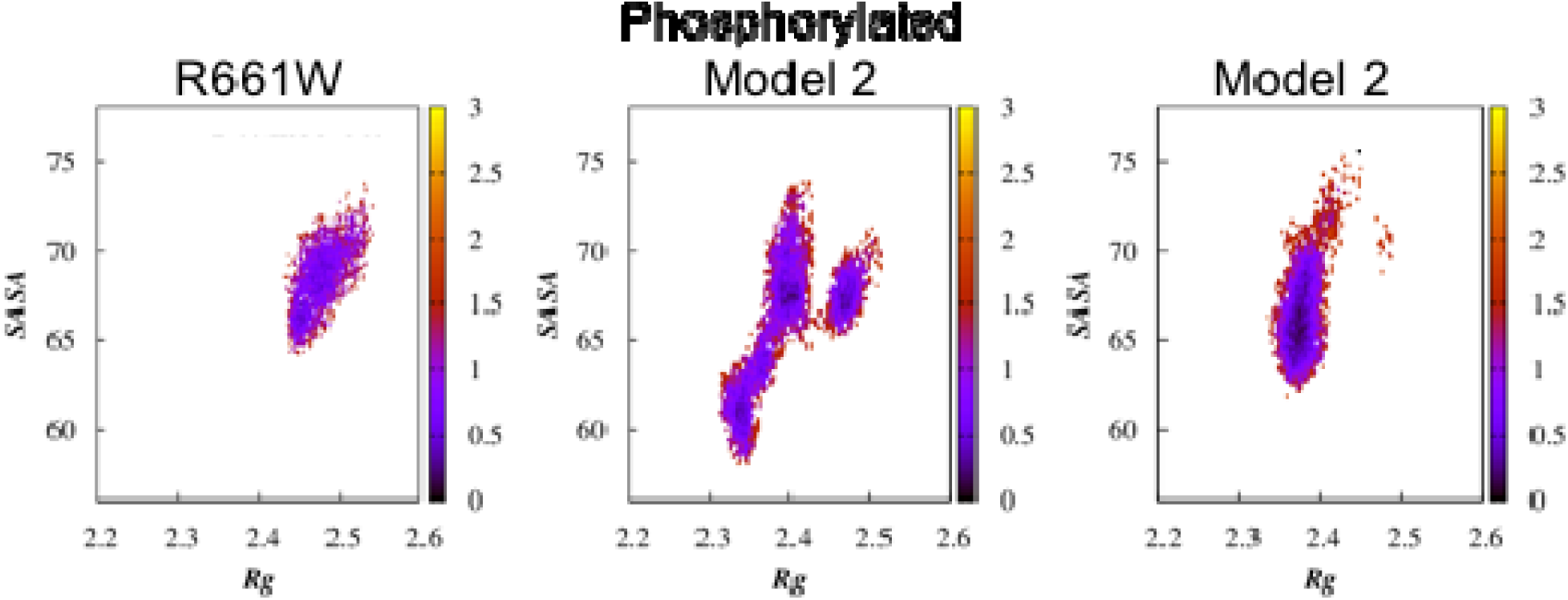
Relationship between the radius of gyration (Rg), solvent-accessible surface area of loop region (SASA) and the surface free energy (ΔG in kcal/mol) change of unphosphorylated and phosphorylated model2, and R661W.

## 4.0 DISCUSSION

We have extensively characterized three computational models of pRbAB with complete pRbPL linker and have studied the mechanism of Ser608 and Ser612 phosphorylation mediated changes in interaction with E2FTA. All the three models, two from ROBETTA and one from AlfaFold, show a short helix formed by residues 602-607 (Fig. 2) that was also observed in a crystal structure PDB:4ELL that contained a shortened helix and a phosphomimetic mutation of Ser608 [21]. The helix in the 4ELL crystal structure mimics a similar helix in the C-terminal of E2FTA, and has been reported to interact with critical residues that are also involved in the interaction of E2FTA with the AB-pocket [21]. While the phosphorylation of Ser608 and Ser612 in pRbPL loop has been proposed to competitively inhibit E2F binding due to formation of an intramolecular interaction of the pRbPL loop with the pocket domains[21, 22], all the three models showed the presence of this short helix with or without phosphorylation of Ser608 and Ser612, that suggest contribution from other factors for the observed effect of Ser608/Ser612 phosphorylation on E2F binding to pRbAB.

Long protein linkers or loops can present multiple conformations due to their dynamic nature. Our data shows considerable motion of the pRbPL linker region. Moreover, within pRbPL, the region containing Ser608 and Ser612 has the highest mobility and surface exposure in all the three models (Fig. 4C, D). These features can ensure a high probability of these residues to be exposed for phosphorylation. However, the motion of the pRbPL linker does not cause any major movements of the pocket domains with respect to each other as shown by interactions between the pocket domains to be mostly conserved compared to those shown for the crystal structures (Fig. 3A). These observations are supported by previous reports that the pocket domains are rigid and do not undergo any major change upon interacting with other binding partners [31].

While the linker region shows interactions with the pocket domains, the crucial interactions shown in 4ELL structure were not displayed by all the models without or with phosphorylation. Therefore, a competitive mechanism displacing the bound E2F upon phosphorylation of Sere608, may be likely, but may not be the dominant mechanism as shown by the movement of the linker and probability of formation of the short helix between residues 602-610 even without phosphorylation. Further, in the presence of the pRbPL linker, it may be difficult for the TA domain of E2F to penetrate deep into the space between the pocket domains due to interferences from the dynamics of the long linker. Indeed, this is what was observed when the E2FTA was docked to the pocket domain connected with the linker (Fig. 6). Our results show that the phosphorylation of Sere608/Ser612 changes the end-to-end distance of the loop. Additionally, phosphate groups can change protein conformation due to electrostatic perturbation. While long-range conformational changes in pRb have been reported in response to phosphorylation of various residues in its linkers [9], the pRbPL phosphorylation does not cause large scale rearrangement of various structural domains of the protein. A significant change in the electrostatic nature of the cavity observed due to phosphorylation of Ser608 and Ser612 in the pRbPL loop was observed (Fig. 7B, H). Additionally, phosphorylation induces a significant change in the compactness and surface free energy change compared to the unphosphorylated pRbAB (Fig 9). Thus, a change in the energy landscape, and electrostatic effects may be the major contributing factors for reduced interactions and destabilization of the loop, resulting in E2F release.

## 5.0 CONCLUSIONS

In general, in many cancers, deletion mutants of pRb are more prevalent, unlike p53 that has a prevalence of missense mutations. However, there are cancer derived mutations clustered around the pocket domain interface^7^, and six missense mutations in the region including the pRbPL linker in cbioportal cancer databases (www.cbioportal.org/). Three of these mutations are in the pRbPL linker. Of these, Tyr606 is a recurrent hotspot and Arg621 mutation is pathogenic or likely to be pathogenic (www.cbioportal.org/). Another frequent missense mutation, R661W, with low affinity for E2F, has a lower solvent exposed surface area, especially in the loop region (Fig. 8) and displays a broadly dispersed conformational landscape compared to full length unphosphorylated pRbAB indicating dynamic conformational states that may negatively impact E2F binding (Fig. 9). The ability of the generated computational models to predict the known effects of R661W mutation on binding to E2F shows the generated models can be utilized to evaluate other missense mutations in pRb whose effects on pRb function have not been tested. The biological significance of any mutations or single nucleotide polymorphisms close to the residues that interact with the LxCxE motif of the HPV E7 protein has not been investigated. A similar analysis of missense mutations in the region of HPV E7 interaction can be tested for their ability to either enhance or prevent the binding of the E7 protein, that will have biological correlates. It is already known that pRbAB is only marginally stable, at least *in vitro*, with its stability dependent on its interactions with binding partners [14]. In this context, it will be of interest to explore whether missense mutations contribute to pRbAB aggregation and how these mutations might impact its interactions with other proteins and its broader physiological functions.

## Supplementary material

Supplementary Figures S1-S2 and supporting Table S1.

### Data and Software Availability

All the software and data used are available in public domain as referred to in text. The models (coordinates and associated data) are available in the link: <u>Associated Data Link</u>

## Acknowledgements

Financial support from DST-BUILDER (BT/PR3000/INF/22/153/2012) grant to JNU is gratefully acknowledged. The authors thank Dr. Ranja Sarkar and Dr. Pinki Dey for initiating and laying the foundation for this work. KK is grateful for fellowship support from the DST-SERB grant CRG/2023/000636 to AB. RS and PD were supported by DST-BUILDER grant, and DST-SERB: ECR/2016/000188 to AB.

## CRediT authorship contribution statement

**Kavana, K:** Writing – original draft, Investigation, Visualization. **Bhattacherjee, A.:** Supervision, Resources, Funding acquisition. **Ghosh, I.:** Conceptualization, Methodology, Supervision, Formal analysis, Writing – review & editing. **Tiwari, S.:** Conceptualization, Supervision, Formal analysis, Writing – review & editing.

## REFERENCES

1. Hanschen ER, Marriage TN, Ferris PJ, Hamaji T, Toyoda A, Fujiyama A, et al. The Gonium pectorale genome demonstrates co-option of cell cycle regulation during the evolution of multicellularity. Nat Commun. 2016;7:11370. doi: 10.1038/ncomms11370.

2. Dick FA. Structure-function analysis of the retinoblastoma tumor suppressor protein - is the whole a sum of its parts? Cell Div. 2007;2:26. doi: 10.1186/1747-1028-2-26.

3. Velez-Cruz R, Johnson DG. The Retinoblastoma (RB) Tumor Suppressor: Pushing Back against Genome Instability on Multiple Fronts. Int J Mol Sci. 2017;18(8). doi: 10.3390/ijms18081776.

4. Yao Y, Gu X, Xu X, Ge S, Jia R. Novel insights into RB1 mutation. Cancer Lett. 2022;547:215870. doi: 10.1016/j.canlet.2022.215870.

5. Knudsen ES, Knudsen KE. Tailoring to RB: tumour suppressor status and therapeutic response. Nat Rev Cancer. 2008;8(9):714–24. doi: 10.1038/nrc2401.

6. Lee C, Chang JH, Lee HS, Cho Y. Structural basis for the recognition of the E2F transactivation domain by the retinoblastoma tumor suppressor. Genes Dev. 2002;16(24):3199–212. doi: 10.1101/gad.1046102.

7. Lee JO, Russo AA, Pavletich NP. Structure of the retinoblastoma tumour-suppressor pocket domain bound to a peptide from HPV E7. Nature. 1998;391(6670):859-65. doi: 10.1038/36038.

8. Liu X, Clements A, Zhao K, Marmorstein R. Structure of the human Papillomavirus E7 oncoprotein and its mechanism for inactivation of the retinoblastoma tumor suppressor. J Biol Chem. 2006;281(1):578–86. doi: 10.1074/jbc.M508455200.

9. Rubin SM. Deciphering the retinoblastoma protein phosphorylation code. Trends Biochem Sci. 2013;38(1):12–9. doi: 10.1016/j.tibs.2012.10.007.

10. Vormer TL, Hansen JB, Te Riele H. The retinoblastoma protein: multitasking to suppress tumorigenesis. Mol Cell Oncol. 2015;2(1):e968062. doi: 10.4161/23723548.2014.968062.

11. Lees JA, Buchkovich KJ, Marshak DR, Anderson CW, Harlow E. The retinoblastoma protein is phosphorylated on multiple sites by human cdc2. EMBO J. 1991;10(13):4279–90.

12. Classon M, Dyson N. p107 and p130: versatile proteins with interesting pockets. Exp Cell Res. 2001;264(1):135–47. doi: 10.1006/excr.2000.5135.

13. Pye CR, Bray WM, Brown ER, Burke JR, Lokey RS, Rubin SM. A Strategy for Direct Chemical Activation of the Retinoblastoma Protein. ACS Chem Biol. 2016;11(5):1192–7. doi: 10.1021/acschembio.6b00011.

14. Chemes LB, Noval MG, Sanchez IE, de Prat-Gay G. Folding of a cyclin box: linking multitarget binding to marginal stability, oligomerization, and aggregation of the retinoblastoma tumor suppressor AB pocket domain. J Biol Chem. 2013;288(26):18923–38. doi: 10.1074/jbc.M113.467316.

15. Kratzke RA, Otterson GA, Lin AY, Shimizu E, Alexandrova N, Zajac-Kaye M, et al. Functional analysis at the Cys706 residue of the retinoblastoma protein. J Biol Chem. 1992;267(36):25998–6003.

16. Kratzke RA, Otterson GA, Hogg A, Coxon AB, Geradts J, Cowell JK, et al. Partial inactivation of the RB product in a family with incomplete penetrance of familial retinoblastoma and benign retinal tumors. Oncogene. 1994;9(5):1321–6.

17. Ramakrishnan C, Subramanian V, Balamurugan K, Velmurugan D. Molecular dynamics simulations of retinoblastoma protein. J Biomol Struct Dyn. 2013;31(11):1277–92. doi: 10.1080/07391102.2012.732345.

18. Papaleo E, Saladino G, Lambrughi M, Lindorff-Larsen K, Gervasio FL, Nussinov R. The Role of Protein Loops and Linkers in Conformational Dynamics and Allostery. Chem Rev. 2016;116(11):6391–423. doi: 10.1021/acs.chemrev.5b00623.

19. Robinson CR, Sauer RT. Optimizing the stability of single-chain proteins by linker length and composition mutagenesis. Proc Natl Acad Sci U S A. 1998;95(11):5929–34.

20. Anthis NJ, Clore GM. The length of the calmodulin linker determines the extent of transient interdomain association and target affinity. J Am Chem Soc. 2013;135(26):9648–51. doi: 10.1021/ja4051422.

21. Burke JR, Hura GL, Rubin SM. Structures of inactive retinoblastoma protein reveal multiple mechanisms for cell cycle control. Genes Dev. 2012;26(11):1156–66. doi: 10.1101/gad.189837.112.

22. Burke JR, Deshong AJ, Pelton JG, Rubin SM. Phosphorylation-induced conformational changes in the retinoblastoma protein inhibit E2F transactivation domain binding. J Biol Chem. 2010;285(21):16286–93. doi: 10.1074/jbc.M110.108167.

23. The National Center for Biotechnology Information. https://www.ncbi.nlm.nih.gov.

24. Yang J, Anishchenko I, Park H, Peng Z, Ovchinnikov S, Baker D. Improved protein structure prediction using predicted interresidue orientations. Proc Natl Acad Sci U S A. 2020;117(3):1496–503. doi: 10.1073/pnas.1914677117.

25. Raman S, Vernon R, Thompson J, Tyka M, Sadreyev R, Pei J, et al. Structure prediction for CASP8 with all-atom refinement using Rosetta. Proteins. 2009;77 Suppl 9(0 9):89–99. doi: 10.1002/prot.22540.

26. Yang J, Yan R, Roy A, Xu D, Poisson J, Zhang Y. The I-TASSER Suite: protein structure and function prediction. Nat Methods. 2015;12(1):7–8. doi: 10.1038/nmeth.3213.

27. Zhang Y. I-TASSER server for protein 3D structure prediction. BMC Bioinformatics. 2008;9:40. doi: 10.1186/1471-2105-9-40.

28. AlphaFold Protein Structure Database. https://alphafold.ebi.ac.uk.

29. Jumper J, Evans R, Pritzel A, Green T, Figurnov M, Ronneberger O, et al. Highly accurate protein structure prediction with AlphaFold. Nature. 2021;596(7873):583-9. doi: 10.1038/s41586-021-03819-2.

30. Varadi M, Anyango S, Deshpande M, Nair S, Natassia C, Yordanova G, et al. AlphaFold Protein Structure Database: massively expanding the structural coverage of protein-sequence space with high-accuracy models. Nucleic Acids Res. 2022;50(D1):D439–D44. doi: 10.1093/nar/gkab1061.

31. Balog ER, Burke JR, Hura GL, Rubin SM. Crystal structure of the unliganded retinoblastoma protein pocket domain. Proteins. 2011;79(6):2010–4. doi: 10.1002/prot.23007.

32. ProSA-web: https://prosa.services.came.sbg.ac.at/prosa.php.

33. Pronk S, Páll S, Schulz R, Larsson P, Bjelkmar P, Apostolov R, et al. GROMACS 4.5: a high-throughput and highly parallel open source molecular simulation toolkit. Bioinformatics. 2013;29(7):845–54. doi: 10.1093/bioinformatics/btt055.

34. Wiederstein M, Sippl MJ. ProSA-web: interactive web service for the recognition of errors in three-dimensional structures of proteins. Nucleic Acids Res. 2007;35(Web Server issue):W407–10. doi: 10.1093/nar/gkm290.

35. Sippl MJ. Recognition of errors in three-dimensional structures of proteins. Proteins. 1993;17(4):355–62. doi: 10.1002/prot.340170404.

36. PyMOL Molecular Graphics System. https://www.pymol.org/sites/default/files/pymol_0.xml.

37. Laskowski RA, Swindells MB. LigPlot+: multiple ligand-protein interaction diagrams for drug discovery. J Chem Inf Model. 2011;51(10):2778–86. doi: 10.1021/ci200227u.

38. Honorato RV, Trellet ME, Jiménez-García B, Schaarschmidt JJ, Giulini M, Reys V, et al. The HADDOCK2.4 web server for integrative modeling of biomolecular complexes. Nat Protoc. 2024;19(11):3219–41. doi: 10.1038/s41596-024-01011-0.

39. Tian W, Chen C, Lei X, Zhao J, Liang J. CASTp 3.0: computed atlas of surface topography of proteins. Nucleic Acids Res. 2018;46(W1):W363–W7. doi: 10.1093/nar/gky473.

40. Chen CR, Makhatadze GI. ProteinVolume: calculating molecular van der Waals and void volumes in proteins. BMC Bioinformatics. 2015;16(1):101. doi: 10.1186/s12859-015-0531-2.

41. Polyansky AA, Zagrovic B. Protein Electrostatic Properties Predefining the Level of Surface Hydrophobicity Change upon Phosphorylation. J Phys Chem Lett. 2012;3(8):973–6. doi: 10.1021/jz300103p.

42. Otterson GA, Chen W, Coxon AB, Khleif SN, Kaye FJ. Incomplete penetrance of familial retinoblastoma linked to germ-line mutations that result in partial loss of RB function. Proc Natl Acad Sci U S A. 1997;94(22):12036–40.

