## Supplementary figures and tables for "Mechanism of phosphorylation dependent interactions of complete Retinoblastoma-AB pocket domain with its linker"

**Figure S1**

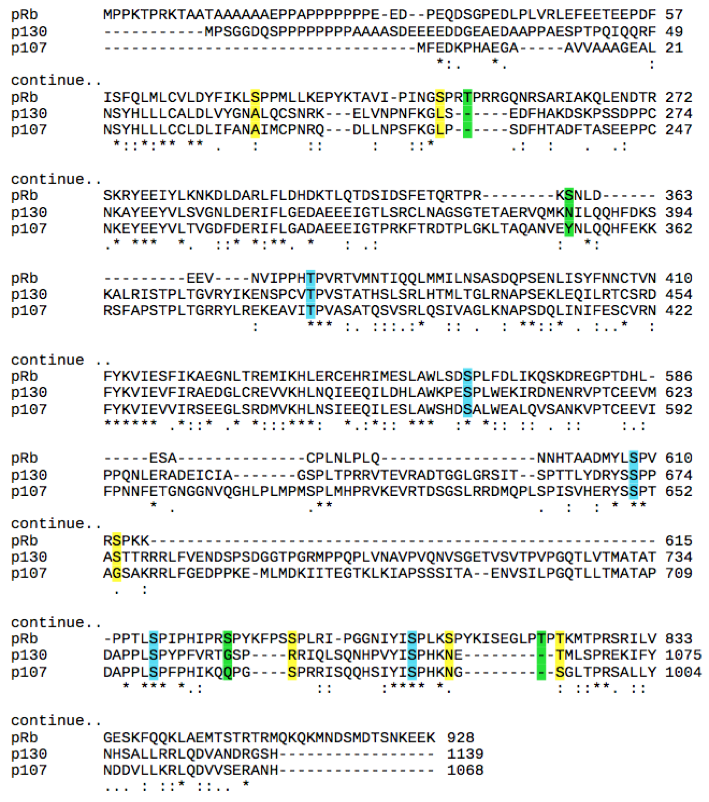

**Figure S1:** Conservation of phosphorylation sites in pRB family proteins. Parts of sequence alignment of pRb homologs are shown to indicate conservation of phosphorylation sites . Blue, conserved in all three homologs; yellow, conserved in two homologs; green, not conserved.

Please add the S/T residue numbers in the figure to show which phosphorylation sites are conserved and otherwise.

**Figure S2**

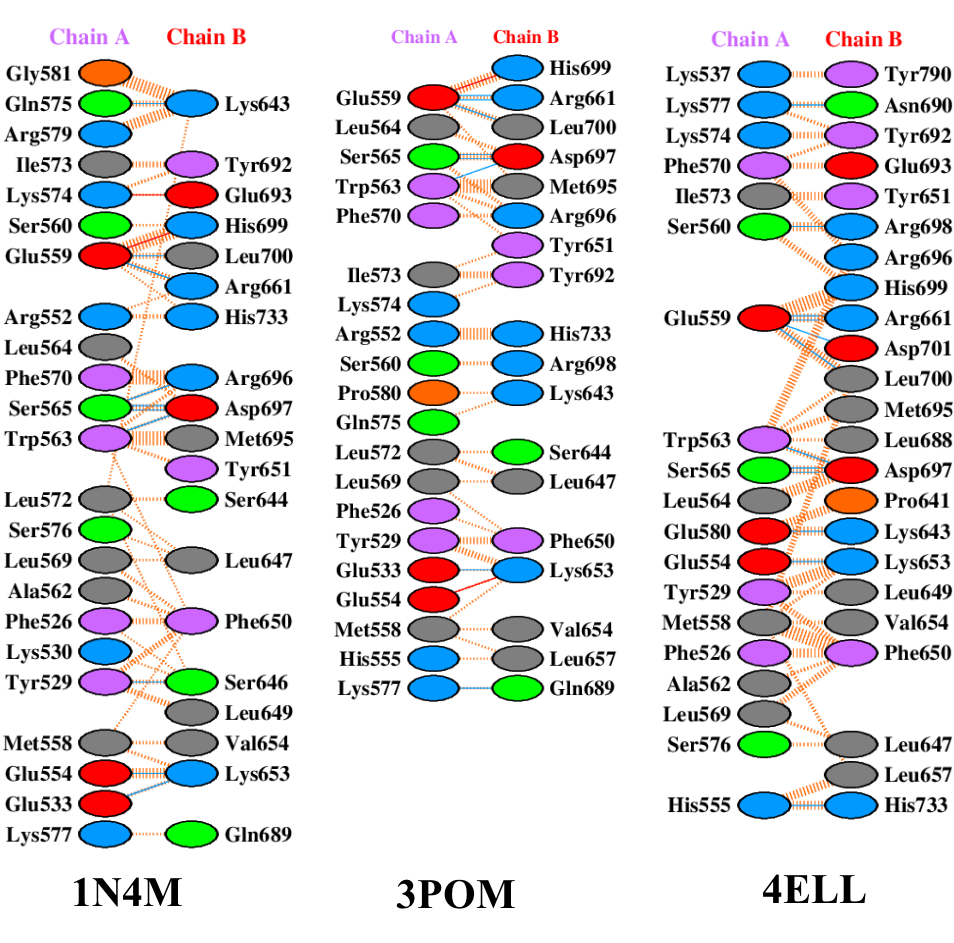

**Figure S2**: Ligplots of the interactions between pocket A and B in the crystal structures 1N4M, 3POM, 4ELL. ChainA, Pocket A; Chain B, Pocket B. Color code for the amino acids is Grey: Neutral; Blue: positively charged; Red: negatively charged. Interactions via the side chains are shown in Green: neutral side chain interactions; Violet: hydrophobic; Yellow: Special cases (Gly,Pro,Cys). Nonbonded interactions are shown with dashed orange and sky blue lines and bonded interactions are shown with continuous red lines.

**Table S1:** The structural analysis of generated models. RMSD against PDB structures 3POM and 1N4M along with Procheck Verify3D and Z-scores are shown.

| Parameters  Models | RMSD(Å)  wrt PDB ID:3POM | | RMSD(Å)  wrt PDB ID: 1N4M | Procheck (R-plot) | | Verify3D | Z-score | PROSA(Z score)  without loop region |
| --- | --- | --- | --- | --- | --- | --- | --- | --- |
| IT-Model1 | 0.852 | | 0.588 | 76.9% | | 87.35% | -8.76 | -8.9 |
| IT-Model2 | 0.928 | | 0.354 | 74.8% | | 89.78% | -7.02 | -8.83 |
| IT-Model3 | 0.921 | | 0.327 | 76.4% | | 93.67% | -8.03 | -8.99 |
| IT-Model4 | 1.399 | | 1.638 | 74.3% | | 87.59% | -7.94 | -8.46 |
| IT-Model5 | 0.842 | | 0.824 | 74.3% | | 94.89% | -9.4 | -8.75 |
| Rob-Model1 | 1.028 | | 1.008 | 85.8% | | 84.91% | -10.47 | -9.33 |
| Rob-Model2 | 1.071 | | 0.972 | 89.0% | | 86.37% | -10.48 | -9.85 |
| Rob-Model3 | 1.014 | | 0.986 | 90.3% | | 85.89% | -9.62 | -9.31 |
| Rob-Model4 | 1.026 | | 0.909 | 91.1% | | 91.48% | -10.33 | -9.81 |
| Rob-Model5 | 1.082 | | 1.009 | 90.3% | | 84.18% | -10.65 | -9.71 |
| Alfa-Model | 0.573 | | 0.610 | 85.8% | | 84.91% | -10.44 | -9.31 |
| Reference PDB structures | | | | |  |  |  |  |
| Models | | PROSA(Z score)  without loop region | | |  |  |  |  |
| 3POM | | -9.31 | | |  |  |  |  |
| 1N4M | | -6.97 | | |  |  |  |  |
| 4ELL | | -8.16 | | |  |  |  |  |
