## Supplementary material for "Mechanism of phosphorylation dependent interactions of complete Retinoblastoma-AB pocket domain with its linker": Table 1

Table 1: Docking scores of the Model2 with the E2FTA under different phosphorylation states.

| **System** | **HADDOCK Score** | **No of Hydrogen bonds** | **Hydrogen bond forming residues (Rb)** | **Hydrogen bond**  **forming  Residues (E2F)** |
| --- | --- | --- | --- | --- |
| Crystal Structure Docked: 1N4M | -98.667 | 10 | ILE785, ARG656, HIS555, GLU551, LYS537, ARG467 | LEU413, ASP411, ASP410, TRP414, GLU420, GLU417, ASP424, SER423, GLY419 |
| Unphosphorylated | -96.779 | 9 | LYS640, SER788, ARG656, HIS600, GLU587, HIS585, GLU533 | ASP424, ASP410, ASP411, SER423, TRP414, GLY415, TYR412 |
| Phosphorylated (S608, S612) | -110.1+/-2.2 | 4 | ARG656, THR645, GLU533 | ASP411, TRP414, ASP410 |
