## Supplementary material for "Mechanism of phosphorylation dependent interactions of complete Retinoblastoma-AB pocket domain with its linker": Table 2

Table 2: Docking scores of the Model1 R661W mutant with the transactivation domain of E2F.

| **System** | **Binding energy(Kcal/mol)** | **Z score** | **Number of hydrogen bonds** | **Hydrogen bond forming residues (Rb)** | **Hydrogen bond**  **forming  Residues (E2F)** |
| --- | --- | --- | --- | --- | --- |
| Model2-R661W | -94.0 +/- 8.7 | -1.7 | 11 | LYS462, HIS555, TYR790, ARG787, ARG656, LYS653, LYS537 | ASP410, ASP411, GLY415, GLU417, TRP414, ASP427, TYR412 |
